# Intermanual transfer of visuomotor adaptation is related to awareness

**DOI:** 10.1101/617407

**Authors:** Susen Werner, Heiko K. Strüder, Opher Donchin

**Author notes:** Corresponding author: Susen Werner, Institute of Movement and Neurosciences, German Sport University, Am Sportpark Müngersdorf 6, 50933 Köln, Germany.

## Abstract

Previous studies compared the effects of gradual and sudden adaptation on intermanual transfer to find out whether transfer depends on awareness of the perturbation. Results from different groups were contradictory. Since results of our own study suggest that awareness depends on perturbation size, we hypothesize that awareness-related intermanual transfer will only appear after adaptation to a large, sudden perturbation but not after adaptation to a small sudden perturbation or a gradual perturbation, large or small. To confirm this, four groups (S30, G30, S75, G75) of subjects performed out-and-back reaching movements with their right arm. In a baseline block, they received veridical visual feedback of hand position. In the subsequent adaptation block, feedback was rotated by 30 deg (S30, G30) or 75 deg (S75, G75). This rotation was either introduced suddenly (S30, S75) or gradually in steps of 3 deg (G30, G75). After the adaptation block, subjects did an awareness test comprising exclusion and inclusion conditions. The experiment concluded with an intermanual transfer block, in which movements were performed with the left arm under rotated feedback, and a washout block again under veridical feedback. We used a hierarchical Bayesian model to estimate individual movement directions and group averages. The movement directions in different conditions were then used to calculate group and individual indexes of adaptation, awareness, unawareness, transfer and washout. Both awareness and transfer were larger in S75 than in other groups, while unawareness and washout were smaller in S75 than in other groups. Furthermore, the size of awareness indices correlated to intermanual transfer across subjects, even when transfer was normalized to final adaptation level. Thus, we show for the first time that the amount of intermanual transfer directly relates to the extent of awareness of the learned perturbation.

## Introduction

There has been an enduring public interest in the topic of two-footedness in world-class soccer players. Regardless of the actual prevalence of true two-footedness in elite players there is widespread agreement that players who can skillfully use both legs are more versatile and can choose between more options for action than players with strong foot dominance. Against this background, transferability of motor learning to different effectors is of strong interest for the field of athletic training. This holds for elite athletes, but it is also true for recreational sports and also has implications for rehabilitation. In rehabilitation, the transfer of motor skills from the trained to the untrained arm can be useful in the therapy of brain-lesioned patients [1] or in upper-limb prosthesis training [2–4].

Intermanual transfer after sensorimotor adaptation has been extensively studied. Sensorimotor adaptation is a specific type of motor learning where an existing motor behavior is altered in response to environmental changes [5–7]. One common paradigm has participants perform well known movements like reaching movements to computer generated targets. After initial training the environment is perturbed by means of a force field that pushes on the moving arm (dynamic perturbation) or by changes of visual feedback (visuomotor perturbation). These perturbations drive adaptation so that reaching behavior returns to baseline despite the perturbation. Intermanual transfer is measured by testing performance with the untrained limb after adaptation in the trained limb is complete.

Awareness is thought to be among the factors influencing intermanual transfer. Malfait and Ostry (2004) compared dynamic adaptation under two conditions to directly assess the influence of awareness on transfer: force fields were introduced either suddenly or gradually. Sudden introduction caused large initial movement errors and, presumably, awareness of the perturbation. Gradual introduction did not cause large errors and seemed to be learned without awareness. After adaptation, the untrained left arm showed a benefit only after sudden introduction of the force field. However, in visuomotor adaptation, two groups (using small magnitudes of perturbation) did not see a difference [9,10]. Indeed, even explaining the perturbation to participants (presumably leading to full awareness) did not lead to increased intermanual transfer compared to adaptation without explanation [9]. Nor did providing an explicit strategy lead to an effect [10]. Accordingly, both groups concluded that awareness has a negligible effect on intermanual transfer of visuomotor adaptation.

These contradictory results could be attributed to a different pattern of intermanual transfer following adaptation to a force field or a visuomotor perturbation. Yet, it is also possible that the previous studies on visuomotor adaptation failed to demonstrate the effect of awareness on transfer because the chosen perturbation sizes (32 deg in Wang et al. (2011a) and 22.5 deg in Taylor et al. (2011)) did not cause awareness. Perturbations of 20 deg have been shown to induce almost no awareness, even when explained to subjects in advance [12,13]. In at least one study, even perturbations as large as 40 deg engaged very little awareness [12]. Hence, the different manipulations of adaptation condition may have not resulted in the desired effect on awareness of the perturbation. This is an issue because studies of intermanual transfer, generally, did not test awareness directly. Rather, they either assumed that sudden and gradual introduction of the perturbation would cause differential levels of awareness [8], or else they tested for a difference using a post-session questionnaire [9,10]. It has been argued that verbal responses may not reveal all a subjects’ knowledge because the knowledge is held with low confidence or because of the difference of retrieval contexts (e.g., Eriksen 1960; Nisbett and Wilson 1977; Shanks et al. 2005).

A growing interest in explicit and implicit processes of sensorimotor adaptation has seen the development of methods for assessment of cognitive aspects such as awareness of adaptation. Some methods rely on subjects reporting their intended direction of movement (termed *prediction methods* in cognitive science). This can be done in various ways such as indicating when a rotating line matches their aiming direction [17,18] or reporting the direction explicitly with the help of a circular display of landmarks [19]. Using such methods, Poh et al. (2016) found that visuomotor rotation to 45 deg had both explicit and implicit components. However, since reporting continued during transfer, the paradigm may have encouraged an explicit transfer (as subjects made an effort to maintain consistency with the previously reported strategy). In addition, prediction methods can be based on feelings of familiarity [21] and lead to an overestimation of awareness [22].

Jacoby (1991) introduced the process dissociation procedure (PDP) as a method for determining awareness and it is now in wide use by cognitive scientists (e.g., Destrebecqz and Cleeremans 2001; Karabanov and Ullén 2008). The PDP is based on defining conscious knowledge as controllable knowledge. Thus, aware and unaware learning can be estimated by comparing performance when participants attempt to either express or repress a learned behavior. We recently implemented this method in the field of sensorimotor adaptation [12]. We were able to directly measure awareness and unawareness indices after adaptation to different rotation angels, and to show that awareness depends on perturbation size. Our current study applies the PDP to measure the involvement of awareness in transfer of visuomotor adaptation to the untrained limb. We compare intermanual transfer after sudden and gradual adaptation to different sizes of visual perturbations. We measure the participants’ awareness of the perturbation by means of the PDP. This should address the criticism that previous findings were tainted by systematic reinforcement of explicit strategies. If awareness is involved in intermanual transfer of visuomotor adaptation, we should find the amount of transfer to be correlated with the amount of awareness. Our hypothesis is that we will find intermanual transfer only after sudden adaptation to a large perturbation but not after sudden adaptation to a small perturbation or after gradual adaptation.

## Methods

### Participants

Twenty-four female and twenty-four male subjects participated in the study and were randomly assigned to four groups. Their ages ranged from 18 to 29 years and were all adults in the country of testing. Participants were gender and age-matched between the four groups and all participants were right-handed as confirmed by the Edinburgh Handedness Inventory. None of the subjects had any experience in visuomotor adaptation research or exhibited overt sensorimotor deficits besides corrected vision. The experimental protocol was conducted according to the principles expressed in the Declaration of Helsinki and was pre-approved by the local ethical committee. All subjects gave written informed consent.

### Task

Seated subjects performed reaching movements with a pen on a digitizing tablet (Fig. 1). They watched a computer screen through a mirror, such that the virtual image of the screen appeared in the same plane in which subjects performed their movements. The mirror prevented vision of the arm, and the position of the pen was registered and displayed to the participants in real-time as a light-blue cursor on the screen. A yellow starting dot appeared in the center of the virtual display and was then replaced by one of eight possible yellow target dots for a duration of 1000 ms. Target directions were 0 deg, 45 deg, 90 deg,… and 315 deg. The targets were presented in random order and all dots were 5 mm in diameter. The subjects were asked to point as quickly and accurately as possible between the central starting dot and the target. Movements continued for episodes of 35 s duration and were interrupted by rest breaks of 5 s.

**Fig. 1:**
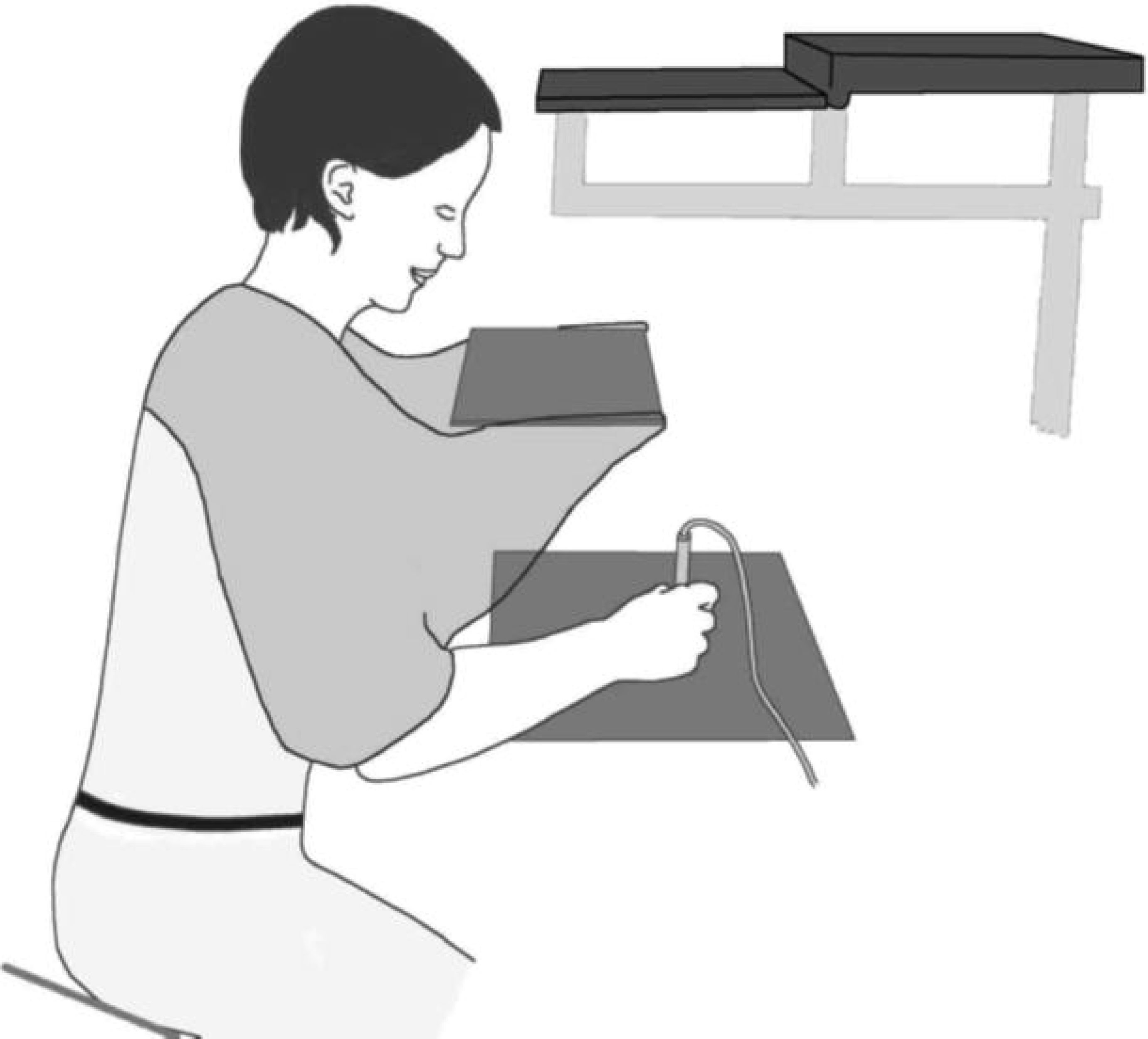
Scheme of experimental apparatus. Shown are tablet (T), screen (S) and mirror (M).

An overview of the experimental protocols is given in Table 1. After *familiarization* with veridical feedback all participants conducted *baseline* episodes without visual feedback, i.e. no cursor visible, as well as baseline episodes with the left and right hand. Depending on group, the perturbation during the following *adaptation* phase was introduced either suddenly or gradually in steps of 3 deg per episode. The first (G30) and second group (S30) were gradually or suddenly exposed to 30 deg rotated visual feedback, whereas the third (G75) and fourth group (S75) were gradually or suddenly exposed to 75 deg rotated visual feedback. Next came two episodes each of *inclusion* and *exclusion* to test for awareness in a process dissociation procedure as in [12,21]. Before inclusion subjects were instructed to ‘use what was learned during adaptation’ and before exclusion subjects were asked to ‘refrain from using what was learned, perform movements as during baseline’ [12]. The order of inclusion and exclusion episodes was randomized between participants. After that, *transfer* to the left hand was tested during two further episodes. No visual feedback was given during inclusion, exclusion and transfer episodes. In particular, visual feedback was not given during transfer episodes in order to prevent confounding transfer with learning benefits to opposite limb learning (“learning to learn”) [20]. Learning was *refreshed* during two intermediate episodes each under rotated feedback. The experimental protocol concluded with a *de-adaptation* phase under veridical feedback.

**Tab. 1:**
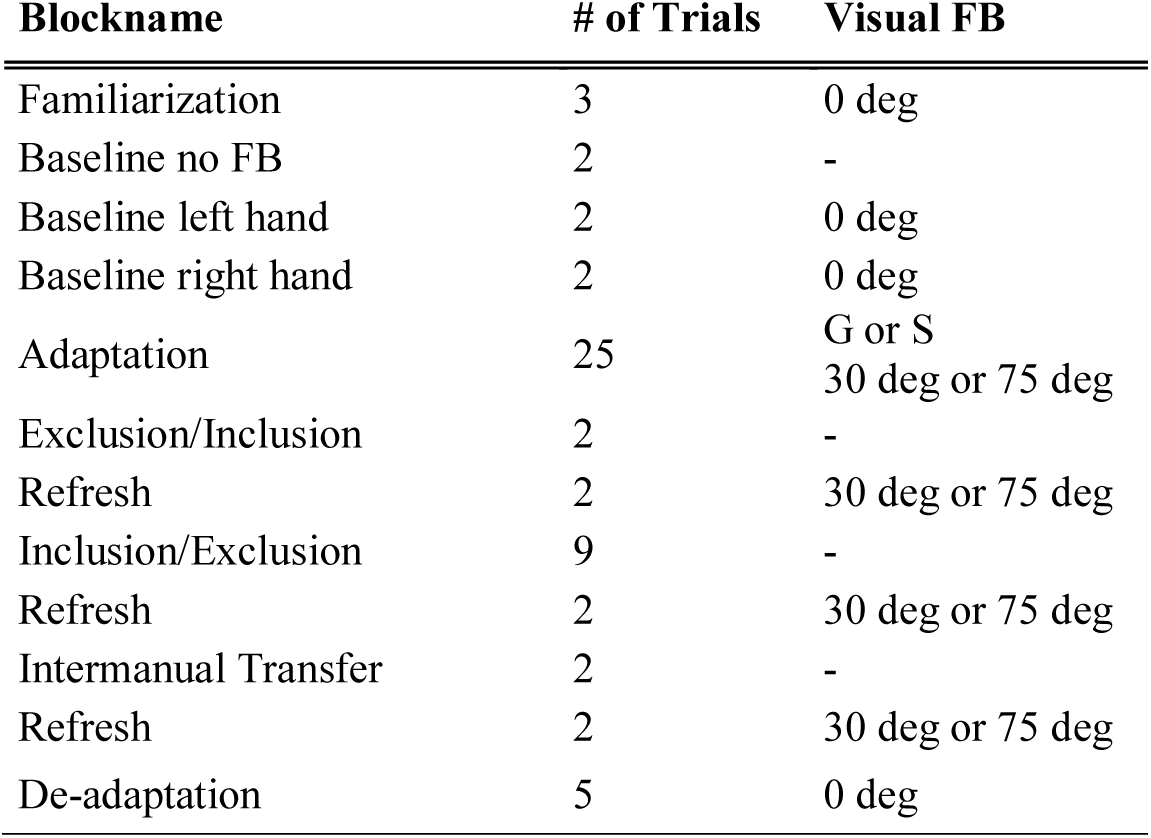
Experimental protocol. Visual feedback (FB) was either not present (-), veridical (0 deg) or rotated (30 deg or 75 deg) gradually (G) or suddenly (S). In the gradual condition rotation size was increased in steps of 3 deg per episode. The order of exclusion and inclusion was alternated between participants.

### Data processing

We quantified participants’ reaching performance as movement direction with respect to the target 150 ms after movement onset, i.e., before feedback-based corrections could become effective [25]. In order to allow the comparison of adaptation to the different rotation sizes, we calculated normalized indices for the different parameters of each participant. Adaptation index, transfer index and washout index were determined as

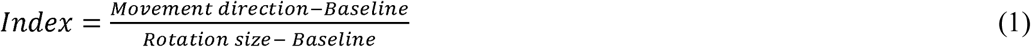

with *Movement direction* equal to mean movement direction of: movements in an adaptation episode for adaptation index for that episode; movements in both transfer episodes for transfer index; movements in the first two washout episodes for washout index. *Baseline* was the mean movement direction of all baseline episodes of the right hand for adaptation and washout index and of the left hand for transfer index. Note, that we defined the amount of transfer as absolute values of transfer episodes as done previously (e.g., Sainburg and Wang 2002).

To account for differences of the amount of adaptation we also calculated a normalized transfer index as

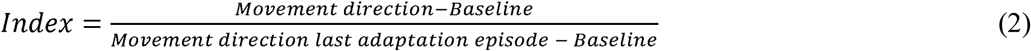

with *Movement direction last adaptation episode equal* to the mean direction of movements in the last adaptation episode. *Movement direction* and *Baseline* were again determined as above.

We further determined awareness and unawareness from reaching performance during exclusion and inclusion episodes. To this end we first calculated exclusion and inclusion indices as

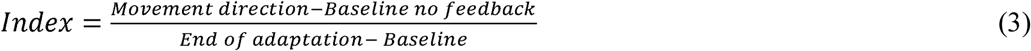

Here, *Movement direction* was the mean movement direction of exclusion or inclusion episodes for the exclusion and inclusion indices, respectively, and *End of adaptation* is taken from the last episode of adaptation phase. According to the PDP, an estimate of awareness can be obtained from the difference between exclusion and inclusion performance and an estimate of unawareness can be obtained from the difference between exclusion performance and baseline. Thus, we calculated the awareness index as inclusion minus exclusion index and the unawareness index equals the exclusion index as in our previous work [12].

We performed a Bayesian statistical analysis. Data and code for this analysis is provided on OSF (https://osf.io/d6swf/). The Bayesian approach samples the joint probability distribution of the parameters given the data [27] instead of focusing on achieving a single best possible estimate and estimating its variance. Our approach followed closely that described in [27]. We fit a linear model to the mean movement directions of each subject in each episode. The linear model had a term for the episode type (34 types for 48 episodes: baseline no feedback, baseline left, baseline right, 25 adaptation episodes, exclusion, inclusion, transfer, refresh, early washout, and late washout), subject group (S30, G30, S75, and G75), and subject (see supporting information S1). It also included terms for the interaction of episode and group and episode and subject. All the coefficients of the linear model were assumed to be normally distributed with a broad, uninformative prior. Noise around the linear model was assumed to have a normal distribution with a standard deviation that varied with subject. The per-subject standard deviation had a gamma distribution whose parameters were sampled hierarchically and had broad priors.

We sampled the posterior distribution of this model given our data using JAGS (4.2.0, http://mcmc-jags.sourceforge.net/) called from Matlab using *matjags* (http://psiexp.ss.uci.edu/research/programs_data/jags/). We used 4 chains, 1,000 adaptation samples, 4,000 burn in samples and 20,000 samples per chain on which we performed our analysis. Chains were initialized using the bootstrap approach described in [27]. We analyzed the standard diagnostics described there as well to ensure that chains had converged for all parameters, that the results were consistent across chains, and that the number of effective samples was in the range of 16,000-20,000 for variance parameters and in the range 22,000- 70,000 for location parameters.

Using the sampled estimates of the linear coefficients, we reconstructed the posterior distribution of the mean movement direction for each episode type for each subject and for each group. We then used these means to calculate samples from the posterior distribution of the indices described above: adaptation (for each adaptation episode), awareness, unawareness, transfer and washout.

We present our results for all indices both for individual subjects and for the group primarily using the 95% high density interval (HDI). This is a region containing 95% of the posterior distribution where every point in the region has higher probability than any point outside the region. Furthermore, for group comparisons we specified a *region of perceived equivalence* (ROPE) of −0.05 through 0.05 around a value of 0 difference between the groups.

Probabilities of less than 0.01 % are reported as negligible. We report the percentage of the HDI that lies within the ROPE as a measure of the probability that the values are equivalent. All analyses were performed using Matlab (Version R2018b).

## Results

Figure 2 shows mean angular movement directions of all experimental phases and groups. (Data is provided as supporting information S2). Note that exclusion is depicted here before inclusion although, in fact, their order was randomised between participants. We did not expect any order effects since we did not find any in our previous study [12]. However, in G75 and S75 we found more awareness and more transfer in participants doing first exclusion and then inclusion. The results of this analysis are provided as supporting information (S3 and S4). Movement directions are near zero during baseline (BAS) for all four groups. All groups show adaptation during the adaptation phase (ADAP). The exceptionally large value in the first adaptation episode of S75 is probably due to irregular reaching behavior during this episode in this group (for sample movement paths for each group see supporting information S5). While angular movement directions decrease gradually in group G30 and G75, they become abruptly negative in group S30 and S75. The two groups adapting to a rotation of 30 deg reach a similar level of movement directions after the first half of adaptation phase which is maintained throughout the subsequent phases. The two groups adapting to a rotation of 75 deg are moving to different directions at the end of adaptation: adaptation is nearly complete in S75 and much less so in G75. Mean group movement directions by the last episode of adaptation phase were −25.5 ± 1.0 (G30), −24.5 ± 0.7 (S30), −51.5 ± 2.4 (G75) and −70.3 ± 5.0 (S75). The level of adaptation is maintained in the subsequent refresh phases in G30 and S30 while it seems to increase in G75 and decrease in S75 (R). The inclusion phase (IN) shows essentially the same level of adaptation, as well. During the exclusion phase, the S30 and G30 groups continue to perform similarly, while the S75 and G75 groups have an inversion. Despite larger adaptation in the S75 group, it shows a much greater ability to exclude this learning than the G75 group in the exclusion phase (EX). This effect is repeated in the washout phase (DA), where S75 washes out much more quickly than G75. The groups showed different levels of partial transfer in the transfer set (TL).

**Fig. 2:**
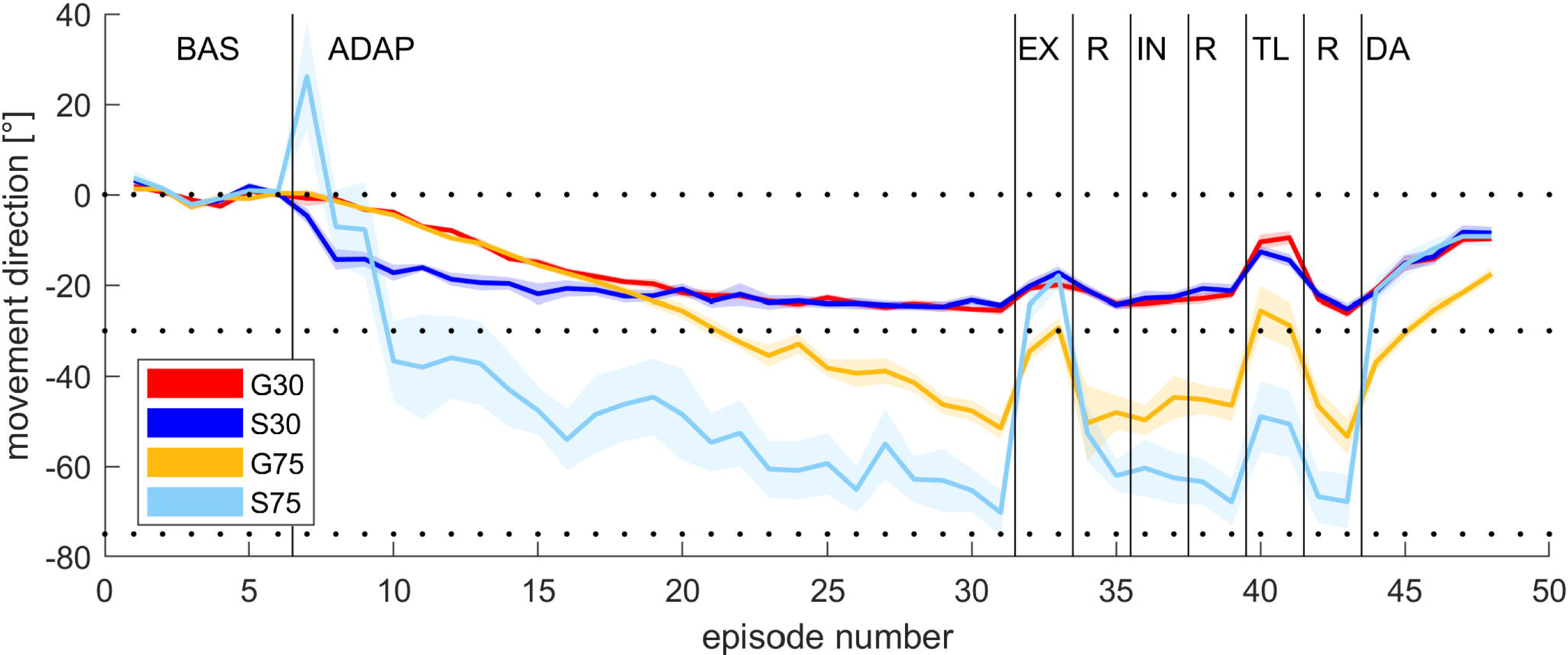
Mean angular movement errors of all blocks and groups. Shown are baseline (BAS), adaptation (ADAP), exclusion (EX), refresh (R), inclusion (IN), intermanual transfer (TL) and de-adaptation (DA). Lines indicate across-subject means, and the shaded area display standard errors.

### Adaptation

To compare learning across groups, we examined the posterior probability distributions for adaptation indices of the group means as well as the posterior probability distributions of the adaptation indexes for individual subjects. Figure 3A depicts the mean of the posterior distribution of the adaptation indices for each group overlaid on its HDI. All groups show adaptation. In addition, adaptation in G75 rises more slowly than G30. This is because adaptation rate in these groups was fixed to 3 deg per movement and so the relative rate in G75 is slower. To compare final adaptation, we calculated the posterior distribution of the difference between the mean of the last 10 episodes for each pair of groups. We stipulated that groups would be considered equivalent if the difference lay in a ROPE of −0.05 to 0.05. Using this procedure, we found a 99.7 % chance that final adaptation of G30 was equivalent to that of S30 and a 91 % chance that it was equivalent to that of S75. Similarly, there was a 94 % chance that final adaptation of S30 and S75 were equivalent. On the other hand, posterior distribution of the difference of final adaptation between all these groups and the G75 group did not overlap at all with the ROPE, indicating less than 0.1 % chance that their relative adaptation is similar to that of G75.

**Fig. 3:**
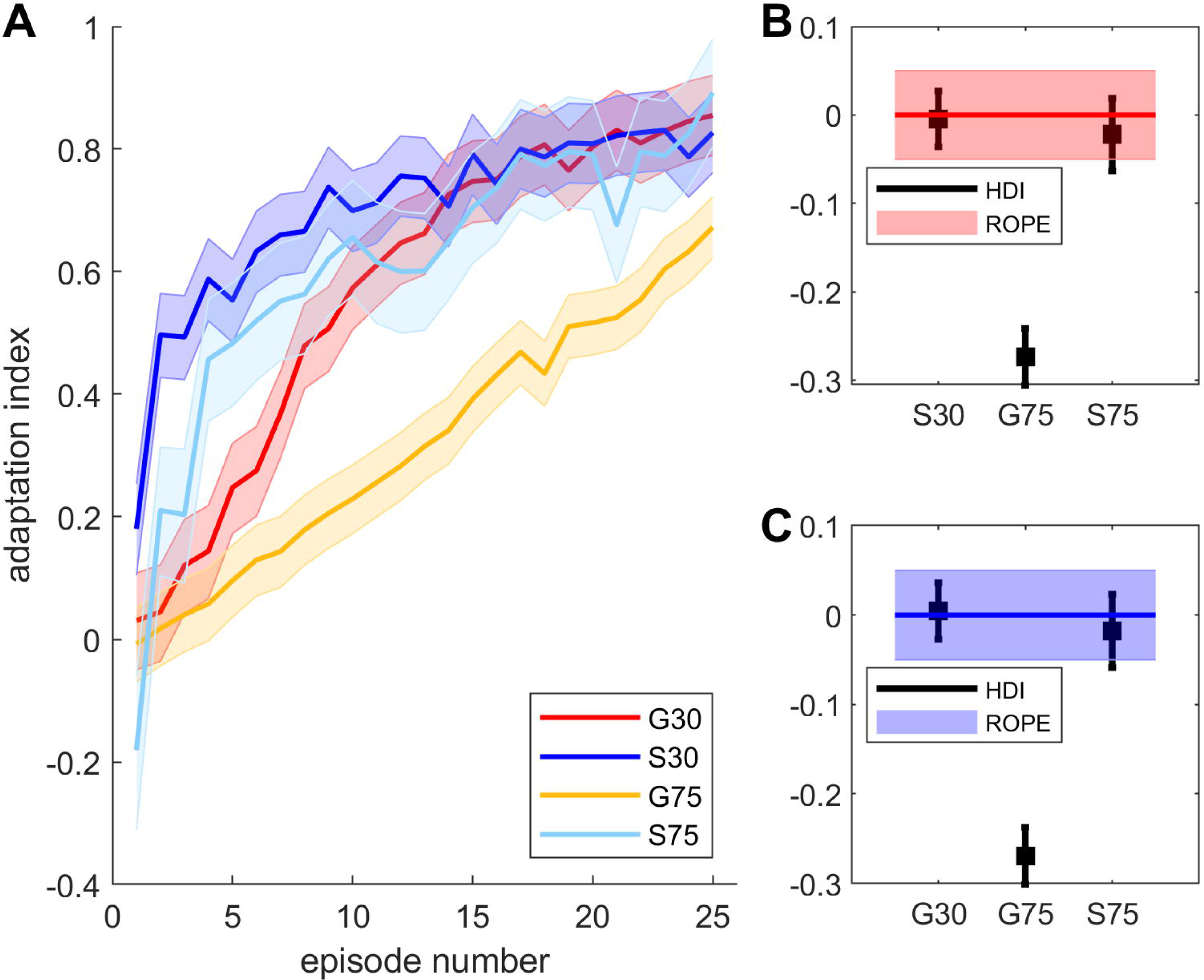
Adaptation. Group means and their HDIs of posterior probability distributions for adaptation indices (A). Average difference from G30 (B) and S30 (C) of the last ten adaptation bins.

These results indicate that learning at the end of adaptation phase was not complete in group G75. Therefore we performed a comparison of the posterior distributions of the difference of the last adaptation episode, all refresh episodes and also the first washout episode within each group. We again determined effective equality using a ROPE of −0.05 to 0.05. For groups G30, S30 and S75 we found a 39 %, 25 % or 39 % chance that final adaptation was equivalent to refresh, respectively. For G75, however, we found a significant increase of index with a chance of equivalence between final adaptation and refresh being less than 0.1 %. Moreover, we compared refresh indices across groups by calculating the posterior distribution of the differences between the estimates of the group indices. This analysis revealed 55 %, 57 % or 54 % chances of equivalence between refresh index of G30, S30 or S75 and that of G75, respectively. These results suggests that learning continued during refresh phase in G75 and final adaptation level approached that of the other groups (for detailed results see supporting information S6).

### Awareness, unawareness, intermanual transfer and washout

Figure 4A shows the means and HDIs of the posterior probability distributions for the group awareness, unawareness, intermanual transfer and washout indices. We performed group comparisons for these indices as was done for the analysis of refresh indices. We show an overview of the results of all group comparisons in Tab. 2A. For S75 we found larger awareness and transfer indices and smaller unawareness and washout indices compared to all other groups. Our analysis revealed less than 0.1 % chance of equivalence for most of these comparisons. While Fig. 4A reveals similar sized indices for G30 and S30, G75 seems to be larger than those two groups in awareness and smaller in unawareness and washout. However, this difference was only significant for washout indices with less than 0.1 % chance of equivalence between G75 and both G30 and S30. Even though Figure 4A suggests a larger transfer in S30 than G30, this difference was not significant with a 3 % chance of equivalence. Thus, statistical analyses reveal a similar pattern of results for awareness and transfer indices with S75 being larger than the other groups and, similarly, the analyses of unawareness and washout indices yield differences of S75 and G75 (for washout only) to all other groups.

**Fig. 4:**
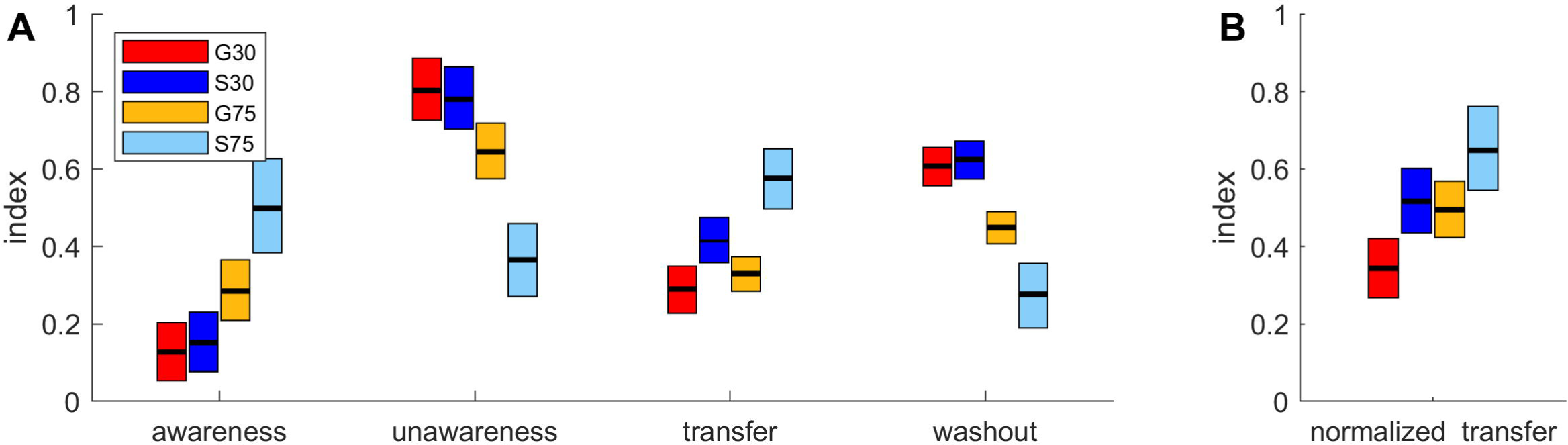
Awareness, unawareness, intermanual transfer and washout. Group means and their HDIs of posterior probability distributions for awareness, unawareness, intermanual transfer and washout indices (A) as well as for normalized intermanual transfer (B).

One consideration is that our indices measure transfer on an absolute scale instead of normalizing for the amount of overall learning. We decided to recalculate the amount of intermanual transfer as a proportion of reaching behavior in the last adaptation episode. Means and HDIs of the posterior probability distributions of this normalized transfer index are shown in Fig. 4B and results of Bayesian analysis in Tab. 2B. The results for normalized transfer resemble those of absolute transfer with regard to a larger index in S75 than in the other groups. However, this difference was only significant (less than 0.1 % chance of equivalence) between S75 and G30. Again we found no significant difference between the two groups adapting to a small rotation with a 2 % chance that normalized transfer index of S30 was equivalent to that of G30.

Figure 5 gives a clearer view of individual subjects performance for awareness, unawareness, intermanual transfer and washout indices. Here each line represents one subject and the range of the line is the HDI for this individual. The subjects are sorted by their mean score on that parameter and colored according to their group. Subjects in the S75 group are, on the whole, in the upper third of the awareness index across subjects and similarly sorted to the top end of the transfer and normalized transfer distributions. It can be seen that there are no G30 or S30 subjects in the top of the population in terms of awareness and no G30 subjects past the median in transfer. Moreover, for the unawareness index the four adaptation conditions led to even clearer group selectivity with the subjects in S75 being (with one exception) the least unaware of all subjects and the subjects in G75 tending to have less unawareness than the other two groups. For the washout index distribution according to groups was also achieved in part.

**Fig. 5:**
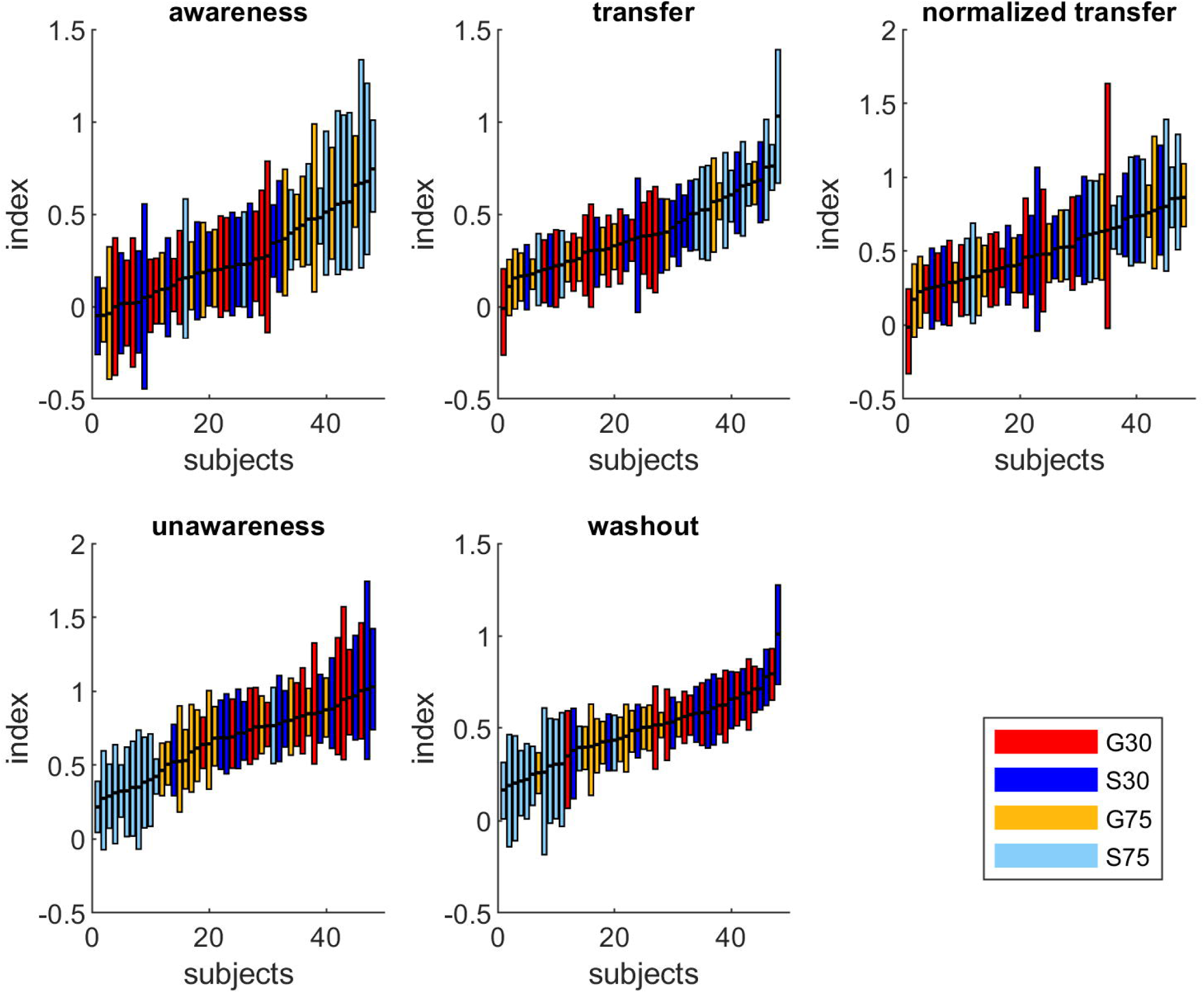
Individual subjects performance for awareness, unawareness, intermanual transfer and washout. Each line represents one subject and the range of the line is the HDI for that individual. Subjects are sorted by their mean score and coloured according to their group.

The results of group means reveal similar patterns for awareness and intermanual transfer indices. Moreover the results of individual subjects performance of these indices reveal considerable inter-individual differences within groups. To further scrutinize the relation between awareness and intermanual transfer we thus calculated correlations between both parameters within and across all groups. Figure 6 demonstrates that larger awareness of the learned perturbation is related to larger transfer (Fig. 6a) as well as to larger normalized transfer (Fig. 6b) to the other limb. The HDI for regression coefficients is 0.26 < r < 0.63 or 0.19 < r < 0.65 for correlations between awareness and transfer or normalized transfer, respectively. In addition, we calculated the HDIs of regression coefficients within each group. Results are shown on Table 2. Note that for transfer as well as for normalized transfer correlations are stronger for the two groups adapting to a large rotation size.

**Tab. 2:** Results of group comparisons. Effective equality of posterior distribution of the differences between the estimates of the four group indices of awareness, unawareness, transfer and washout indices (A) and of normalized transfer index (B) using a ROPE of −0.05 to 0.05 are shown. Significant differences were marked in bold numbers.

| **A** | **Awareness** | **Unawareness** | **Transfer** | **Washout** | **B** | **Norm. transfer** |
| --- | --- | --- | --- | --- | --- | --- |
|  | S75 | S75 | S75 | S75 |  | S75 |
| G30 | < **0.1** % | < **0.1** % | < **0.1** % | < **0.1** % |  | < **0.1** % |
| S30 | < **0.1** % | < **0.1** % | 2 % | < **0.1** % |  | 11 % |
| G75 | 1 % | < **0.1** % | < **0.1** % | 0.4 % |  | 5 % |
|  | G75 | G75 | G75 | G75 |  | G75 |
| G30 | 3 % | 2 % | 59 % | < **0.1** % |  | 3 % |
| S30 | 7 % | 6 % | 16 % | < **0.1** % |  | 60 % |
|  | S30 | S30 | S30 | S30 |  | S30 |
| G30 | 60 % | 58 % | 3 % | 79 % |  | 2 % |

**Tab. 3:** Results of correlations between awareness and intermanual transfer. HDIs of regression coefficients for correlations between awareness and transfer or normalized transfer over all groups and within each group are shown.

|  | **Transfer** | **Normalized transfer** |
| --- | --- | --- |
| All groups | $0.26 < r < 0.63$ | $0.19 < r < 0.65$ |
| G30 | $-0.62 < r < 0.49$ | $-0.60 < r < 0.69$ |
| S30 | $-0.28 < r < 0.66$ | $-0.32 < r < 0.71$ |
| G75 | $0.03 < r < 0.70$ | $-0.05 < r < 0.73$ |
| S75 | $0.06 < r < 0.85$ | $0.17 < r < 0.89$ |

**Fig. 6:**
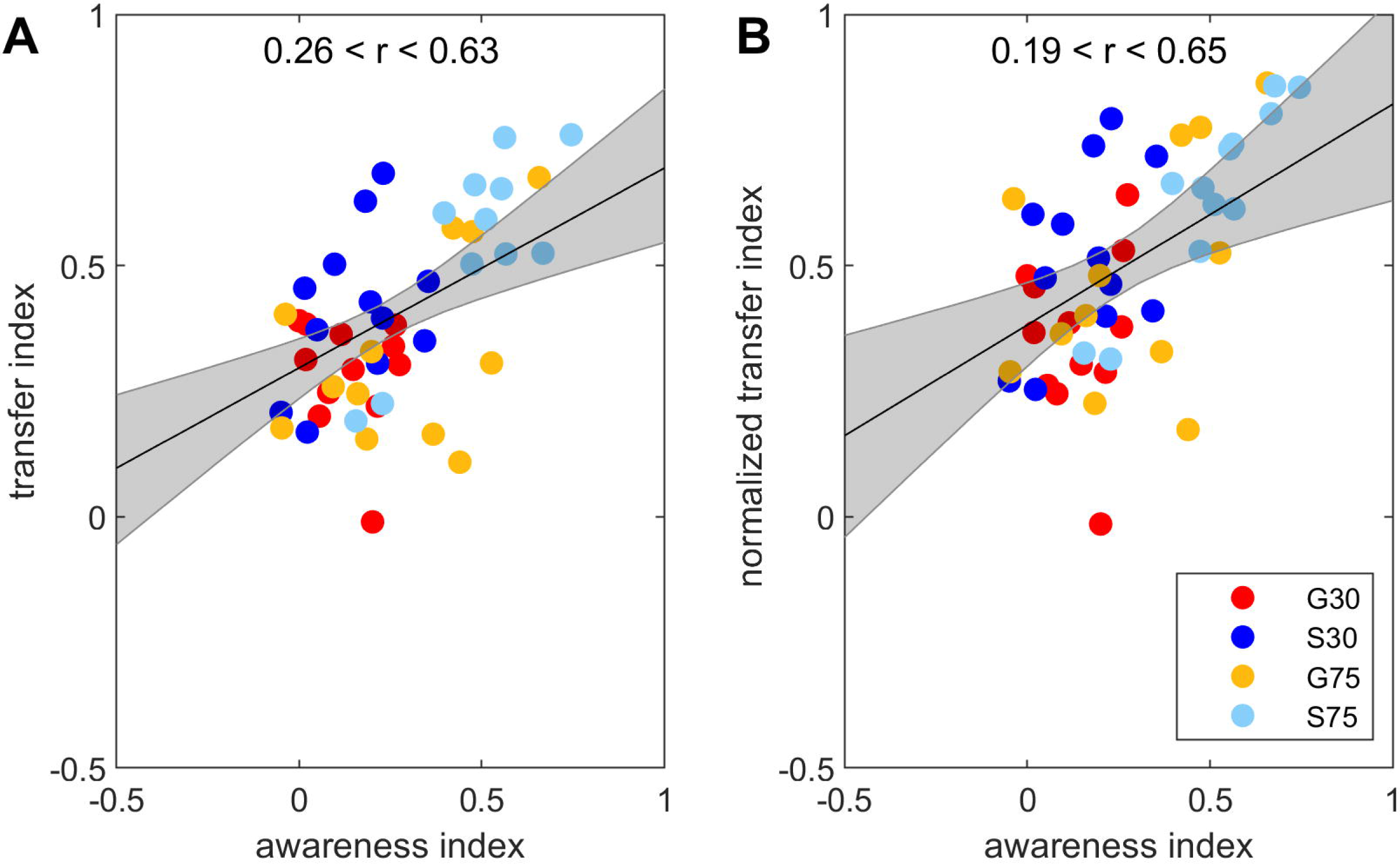
Correlations between awareness and intermanual transfer (A) or normalized intermanual transfer (B). Correlations between the awareness and transfer indices for all participants.

## Discussion

One essential question in sensorimotor adaptation research concerns the generalization or transfer of sensorimotor adaptation: The investigation of transfer to different body segments, different movements or different work-spaces can give insight into the nature of motor learning and the brain structures involved. The aim of the present study was to find out whether the amount of intermanual transfer depends on the degree of awareness of the nature of the perturbation. Therefore, four groups (S30, G30, S75, G75) of participants adapted to a visuomotor perturbation. Visual feedback was rotated either by 30 deg (S30, G30) or 75 deg (S75, G75) and this rotation was either introduced suddenly (S30, S75) or gradually in steps of 3 deg (G30, G75). We also measured awareness and unawareness indices after adaptation with a process dissociation procedure and, additionally, the amount of intermanual transfer to the untrained, non-dominant arm and washout performance. Groups with most awareness seemed to show the most transfer. Awareness was larger in S75 than in all other groups and transfer was larger in S75 than in G75 and G30. This pattern held for both the absolute level of transfer and also, although more weakly, when transfer was normalized by the amount of final adaptation. Furthermore, at the individual level, subjects with more awareness showed more transfer.

### Intermanual transfer

We found intermanual transfer to be more pronounced after sudden adaptation to a large perturbation than after gradual adaptation to either a large or small perturbation. Moreover, we found a strong correlation between awareness and intermanual transfer over all groups, even when transfer was normalized to final adaptation level. These findings support our hypothesis that awareness is indeed involved in intermanual transfer of visuomotor adaptation. Specifically, the relation between the amount of awareness and of transfer was more pronounced after adaptation to large perturbations. Poh et al. (2016) also revealed a contribution of the explicit process of sensorimotor adaptation to intermanual transfer. However, their reporting method of testing explicit adaptation might have actually caused transfer of explicit learning itself as outlined in the introduction. Thus, using the PDP for measuring awareness we could, for the first time, show the involvement of awareness in intermanual transfer of visuomotor adaptation without creating a situation where testing for awareness might reinforce explicit strategies.

Our results also showed an interesting interaction: abrupt introduction of the perturbation led to more transfer when the perturbation was large but not when it was small. This may explain the apparent contradictions in previous studies, with some showing different intermanual transfer after gradual and sudden adaptation [1,8] but others do not [9,10]. Our findings suggest that the latter did not see an effect of awareness on transfer because small perturbation sizes did not induce awareness even in abrupt perturbations. Even explicit instructions did not lead to increased intermanual transfer in those studies [9,10]. It could be argued, that instructing the participants about the nature of the perturbation should lead to awareness. However, awareness of small visuomotor rotations remains minor (about 25% after 20 deg rotation) even after explicit instructions [12].

The amount of intermanual transfer cannot be fully explained by the amount of awareness in the present study. Even after gradual adaptation to a 30 deg rotation with awareness of about 15 % transfer reaches about 30 % and normalized transfer reaches about 35 %. Moreover, while there is no doubt that awareness and transfer are correlated across individuals, these correlations are not strong. The HDI for the correlation coefficient was 0.26 < r < 0.63 and 0.19 < r < 0.65 for transfer and for normalized transfer, respectively. Thus, it is likely that the implicit process of adaptation is also involved in transfer to the other hand as previously suggested [20]. It is equally possible that other factors are involved such as the degree of handedness [28,29].

It is also conceivable that the amount of intermanual transfer in the present study is confounded by group differences of time spend at plateau during learning [30]. While most groups in our study reach a similar plateau early in adaptation and remain there, the G75 does not reach a plateau at all and has less adaptation than the other groups for the last ten episodes. It has been suggested that extended training should lead to greater transfer [30] although these results have not been consistent in the literature [31]. However, because our study included groups with prolonged training at the plateau that did not show strong transfer (the S30 and G30 groups), we do not believe that the result in the G75 group is strongly affected by failing to reach plateau. Thus, we suggest that awareness and not time spent at plateau drive the amount of intermanual transfer.

In order to further confirm this, we tested specifically for normalized transfer in addition to absolute transfer. Even though our results are not as strong for normalized transfer, the general pattern of the results is the same: awareness correlated to both our transfer measures.

### Awareness and unawareness

The degree of awareness of the nature of the perturbation did clearly depend on the perturbations’ magnitude during sudden adaptation. Hence awareness is driven by the size of target error, i.e. the perceived error between cursor and target. This result is consistent with our previous work showing lager awareness after adaptation to a 60 deg than to a 20 deg rotation of visual feedback [12]. For the first time, the degree of awareness was actually measured after adaptation to a gradually introduced perturbation. As suggested earlier [12,32–34] we indeed find very low awareness but large unawareness indices after gradual adaptation.

Performance in aftereffect tests such as pointing directions at the beginning of a washout block is thought to reflect the actual recalibration of sensory-to-motor transformation rules [35,36]. Compared to sudden adaptation, performance in aftereffect tests was previously shown to be improved after gradual saccade [37] and prism adaptation [1], after gradual adaptation to a visual gain [38] and to a 90 deg rotation [39]. Adaptation to smaller visual rotations (30 deg and 60 deg), however, did not lead to a difference of aftereffects between gradual or sudden adaptation [34,40]. This is in line with the present findings. Our analyses reveal a difference of washout indices after gradual or sudden adaptation to a 75 deg but not after gradual or sudden adaptation to a 30 deg rotation. In addition, we find a similar pattern of results for washout and for unawareness indices. Therefore, we can conclude that the amount of recalibration does not depend on the learning condition such as gradual versus sudden, but is negatively related to the amount of awareness gained during adaptation.

It can be argued that gaining awareness does not directly indicate the explicit process of adaptation since it does not ensure the use of explicit strategies [41]. In that respect the irregular use of the term awareness should first be mentioned. Some authors relate to awareness of a perturbation, i.e. the notion that something has changed [11,33], while we specifically refer to awareness of the nature of the perturbation as also done in previous other work [9,42].

Second, the mental construct of awareness might very well be more than just the participants’ report of planned movement directions and the PDP measure and the trial-by-trial reporting might reflect two distinct components of the explicit process. Then again, characterization of implicit and explicit processes over a range of task conditions lead to similar results independent of which method was used for measuring the two processes: The explicit process increases with increasing rotation size and the implicit process takes time to develop during adaptation measured by reporting [19,43] and, likewise, by PDP [12,13]. Further research should be conducted to directly compare the explicit process of sensorimotor adaptation measured by prediction task methods such as reporting of aiming directions [19] or by the process dissociation procedure [12].

### Neuronal correlates of intermanual transfer

It has been suggested that intermanual transfer of sensorimotor adaptation may involve several aspects of a neural representation, some of which are effector dependent and others are effector independent [44]. It can be speculated that cognitive processes such as the explicit process or awareness of a perturbation do not directly relate to a motor action of a specific limb and that they might, thus, relate to an effector independent neural representation. It has been suggested that the explicit processes plays a role in an early stage of the adaptation process [42]. Since several imaging studies on sensorimotor adaptation found early learning to engage prefrontal brain regions (e.g., Clower et al. 1996; Anguera et al. 2007; Inoue et al. 2015), especially the dorsolateral prefrontal cortex [48] this brain area might be related to the explicit process of adaptation. Considering our present findings of intermanual transfer being related to awareness this area may also be related to the transfer of adaptation.

Imaging of intermanual transfer of visuomotor adaptation previously revealed activation in the temporal cortex, the middle occipital gyrus and the right medial frontal gyrus [47]. However, adaptation to a 30 deg rotation was used in that study, which is largely related to the unaware, implicit process of adaptation as suggested previously [12,13] and confirmed by the present data. Accordingly, Anguera et al. (2007) observed no areas of overlap comparing activation at transfer and early adaptation but partial overlap at transfer and late adaptation.

These findings suggest, in combination with our data, that the above mentioned areas might be more related to the transfer of the implicit process of adaptation. Future research should be conducted to scrutinize whether brain areas related to the unaware or implicit process of adaptation are also related to the unaware or implicit process of intermanual transfer. For example, indirect evidence from patient studies supports the involvement of the cerebellum mainly in the implicit process during adaptation itself. For instance, cerebellar patients show decreased aftereffects (e.g., Morton and Bastian 2004; Smith and Shadmehr 2005; Werner et al. 2010; Donchin et al. 2012) or impaired gradual adaptation associated with less awareness [53,54]. However, transcranial direct current stimulation on the trained or untrained hemisphere of the right cerebellum caused faster adaptation, but did not affect intermanual transfer [30].

## Conclusions

In conclusion, the results of this study show for the first time that the size of intermanual transfer directly relates to the extent of awareness of the learned visuomotor perturbation. Our findings can further help explain the disagreement regarding the amount of intermanual transfer after gradual and sudden adaptation by proposing that some studies chose perturbation sizes which left participants adapting to the suddenly introduced distortion just as unaware as those in the gradual group. The amount of transfer cannot be fully explained by awareness in the present study suggesting the involvement of additional factors like the implicit process of adaptation or handedness. Moreover, our findings confirm the idea that participants adapting to gradually introduced perturbations are usually not aware and that the size of aftereffects is negatively related to the amount of awareness gained during adaptation. Our results also underscore the complexity of the implicit and explicit processes of sensorimotor adaptation and emphasize the importance of directly comparing the different tasks of measuring the explicit process or awareness. To sum up, our results can further help explaining the effect of awareness on sensorimotor adaptation and its transfer to the other limb.

## Supporting information

Supplemental Model

Supplemental Results 1

Supplemental Results 2

Supplemental Results 3

## Acknowledgments

This work was supported by a DAAD Travel Grant 91721115 awarded to Susen Werner. The authors thank Mats Harmuth for his help with data collection.

## Supporting information captions

**S1: Model.** Model used for the Bayesian analysis.

**S2: Dataset.** Mean movement directions of each episode (baseline, adaptation, awareness test, refresh, intermanual transfer and deadaptation) for each participant of each group (G30, S30, G75, S75).

**S3: Dataset.** Order of exclusion and inclusion of awareness test for each subject; 1 means exclusion first and 0 means inclusion first.

**S4: Results.** Results of the analysis of the order effect of exclusion and inclusion phase.

**S5: Results.** Original registrations of movement paths of the first adaptation episode produced by a typical participant of each group

**S6: Results.** Results of the analysis of the refresh phase.

