## Supplemental Model for "Intermanual transfer of visuomotor adaptation is related to awareness"

**Model of Bayesian analysis**

```
model{


# N observations (subjects * withinX1 * betweenX2 in my case)
for(i in 1:N){
	# y[i] ~ dt(muY[i], tauY[id[i]], nuY)
	y[i] ~ dnorm(muY[i], tauY[id[i]])
   muY[i] <- a0 + a1[X1[i]] + a2[X2[i]] + a12[X1[i],X2[i]] + aS[id[i]] + a1S[X1[i],id[i]]
}


# Priors


a0 ~ dnorm(0, a0Tau)
for (j1 in 1:nX1) {
	a1[j1] ~ dnorm(0, a1Tau)
}


for (j2 in 1:nX2) {
	a2[j2] ~ dnorm(0, a2Tau)
}
for (j1 in 1:nX1) {
	for (j2 in 1:nX2) {
		a12[j1,j2] ~ dnorm(0, a12Tau)
	}
}
```

```
for(i in 1:nS){
        aS[i] ~ dnorm(0, tauS)
   tauY[i] ~ dgamma(tauYRa, tauYSh)
}

for (j1 in 1:nX1) {
   for (s in 1:nS) {
      a1S[j1,s] ~ dnorm(0, tauA1S)
   }
}

tauYRa <- (tauYMode + sqrt(tauYMode^2 + 4*tauYVar))/(2*tauYVar)
tauYSh <- 1 + tauYMode*tauYRa

tauYMode ~ dgamma(tauYModeSh, tauYModeRa)
tauYVar ~ dgamma(tauYVarSh, tauYVarRa)


# tauS ~ dgamma(tauSRa, tauSSh)
# tauA1S ~ dgamma(tauA1SRa, tauA1SSh)

}
```
