## Supplemental Results 1 for "Intermanual transfer of visuomotor adaptation is related to awareness"

### Results of the analysis of the order effect of exclusion and inclusion phase

Doing exclusion first leads to more awareness and more transfer in the groups doing the 75 degree rotation.

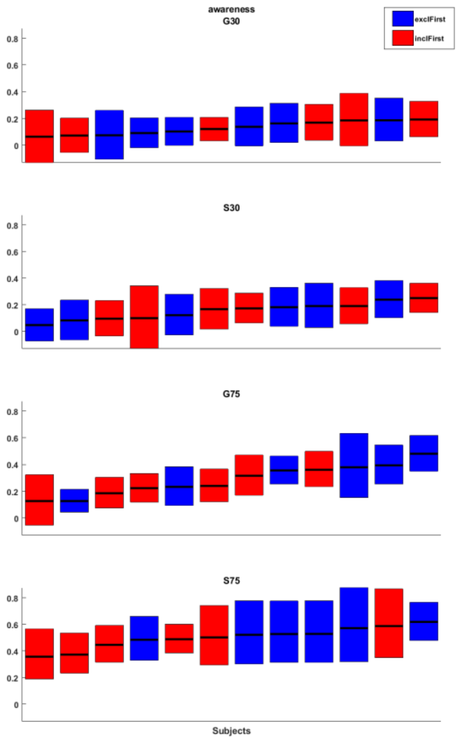


**Fig. 1: Individual subjects performance for awareness.** Each line represents one subject and the range of the line is the HDI for that individual. Subjects who experience exclusion before inclusion are marked in blue, the others in red. All subjects are sorted by their mean score within each group (G30, S30, G75 and S75).

| | G30 | S30 | G75 | S75 |
|---|---|---|---|---|
| hdiLow | -0.0695 | -0.0721 | -0.1510 | -0.1965 |
| mean | 0.0058 | 0.0049 | -0.0749 | -0.0979 |
| hdiHigh | 0.0817 | 0.0810 | -0.0020 | -0.0038 |

**Tab. 1: Results of Bayesian analysis for awareness.** The values of the HDI of the mean difference between exclusion first and inclusion first are shown.

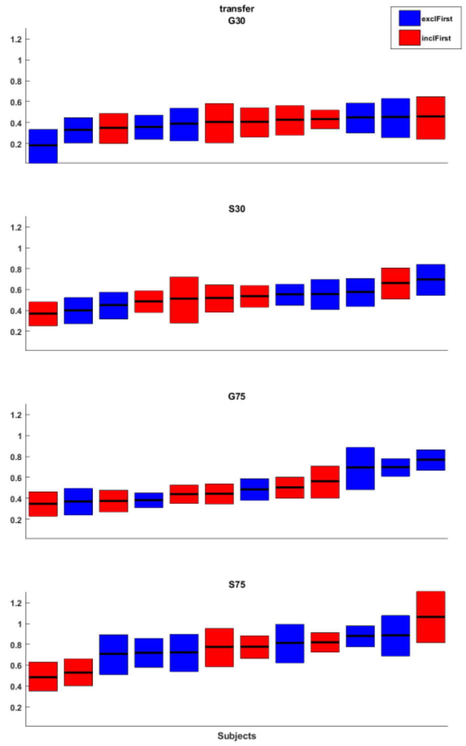


**Fig. 2: Individual subjects performance for transfer.** Each line represents one subject and the range of the line is the HDI for that individual. Subjects who experience exclusion before inclusion are marked in blue, the others in red. All subjects are sorted by their mean score within each group (G30, S30, G75 and S75).

| | **G30** | **S30** | **G75** | **S75** |
|---|---|---|---|---|
| hdiLow | -0.0202 | -0.0579 | -0.1773 | -0.2057 |
| mean | 0.0573 | 0.0117 | -0.1188 | -0.1266 |
| hdiHigh | 0.1350 | 0.0819 | -0.0592 | -0.0484 |

**Tab. 2: Results of Bayesian analysis for transfer.** The values of the HDI of the mean difference between exclusion first and inclusion first are shown.
