## Supplemental Results 2 for "Intermanual transfer of visuomotor adaptation is related to awareness"

### Movement paths of the first adaptation episode of individual subjects

Movement paths in S75 are more irregular than in the other groups.

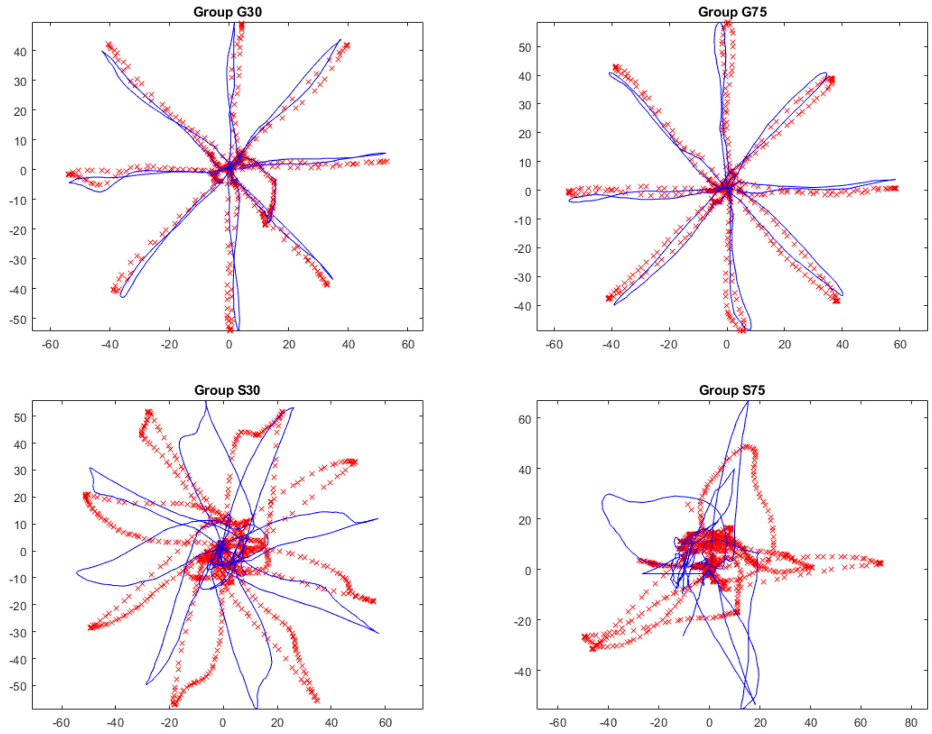


**Fig. 1: Examples of movement paths.** We show original registrations of movement paths (red crosses) and cursor paths (blue lines) of the first adaptation episode produced by a typical participant of each group.
