## Supplemental Results 3 for "Intermanual transfer of visuomotor adaptation is related to awareness"

### Results of the analysis of the refresh phase

Learning continued during refresh phase for group G75.

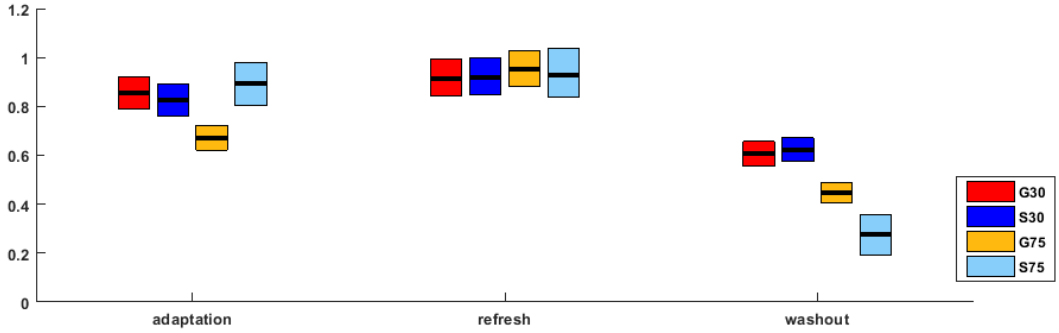


**Fig. 1: Adaptation, refresh and washout.** Group means and their HDIs of posterior probability distributions for the last episode of adaptation, all six refresh episodes and the first washout episode.

| | **Adaptation** | **Refresh** | **Washout** |
|---|---|---|---|
| | S75 | S75 | S75 |
| G30 | 43 % | 56 % | < **0.1 %** |
| S30 | 8 % | 56 % | < **0.1 %** |
| G75 | 8 % | 54 % | < **0.1 %** |
| | G75 | G75 | G75 |
| G30 | 19 % | 55 % | 7 % |
| S30 | 64 % | 57% | 18 % |
| | S30 | S30 | S30 |
| G30 | 19 % | 65 % | 67 % |

**Tab. 1: Results of between group analysis for adaptation, refresh and washout.** Effective equality of posterior distribution of the differences between the estimates of the four group indices of adaptation, refresh and washout indices using a ROPE of -0.05 to 0.05. Significant differences are marked in bold numbers.

| | **G30** | **S30** | **G75** | **S75** |
|---|---|---|---|---|
| | adaptation | adaptation | adaptation | adaptation |
| refresh | 39 % | 25 % | **< 0.1 %** | 39 % |
| washout | **< 0.1 %** | **< 0.1 %** | **< 0.1 %** | **< 0.1 %** |
| | refresh | refresh | refresh | refresh |
| washout | **< 0.1 %** | **< 0.1 %** | **< 0.1 %** | **< 0.1 %** |

**Tab. 2: Tab. 1: Results of within group analysis for adaptation, refresh and washout.** Effective equality of posterior distribution of the differences between the estimates of the adaptation, refresh and washout indices within each group using a ROPE of -0.05 to 0.05. Significant differences are marked in bold numbers.
